# Fever induces long-term synaptic enhancement and protects learning in an accelerated aging model

**DOI:** 10.1101/2025.07.09.664028

**Authors:** Fusheng Du, Qi Wan, Oleg O. Glebov

## Abstract

Physiological impact of fever in the brain remains poorly understood. Here, we demonstrate that induction of fever by yeast injection in rats (N=9) and by whole-body hyperthermia in mice (N=7) triggers structural synaptic enhancement in the prefrontal cortex involving AMPA-type glutamate receptor signalling and protein translation (N=6). Repeated fever induction in juvenile rats (N=9) results in synaptic strengthening that persists into adulthood, mitigating learning deficits and synaptic loss in a D-galactose model of accelerated aging (N=11). Our results show how common environmental conditions may shape brain function in the long-term via synaptic plasticity, warranting further exploration of thermal treatment for cognitive protection in aging.

---

Although increased body temperature represents a common consequence of infection, injury, environmental exposure, and exercise, its physiological consequences remain poorly understood. The brain is particularly susceptible to temperature variations, e.g. even moderate temperature elevation can trigger febrile seizures in children, likely via alteration of neuronal excitability [1, 2]. The longer-term impact of fever on the brain, especially in the context of development and aging, remains unknown.

Our previously published evidence demonstrated that temperature decrease induces synaptic remodeling [3]. To investigate synaptic effects of temperature elevation within the physiological range *in vivo*, we leveraged two well-established rodent models of fever, namely pyrexia by yeast injection in rats [4, 5] and whole-body hyperthermia (WBH) in mice [6], and visualised synapses in the prefrontal cortex (PFC), using immunohistochemistry staining in brain sections to quantify synaptic structure and composition (**Figure S1A**). Both approaches reliably increased body temperature by 2-3°C within 7 hours of treatment (**Figure 1A,B**). Intensity of punctate staining for a synaptic terminal marker Bassoon was increased in both models (**Figure 1C,D**), as was the case for an excitatory postsynaptic density marker Homer (**Figure S1B,C**). Staining intensity for several other synaptic markers was also increased, including Ca_V_2.1, vGlut1, gephyrin, GABRA1, synapsin 1/2, and synaptotagmin 1 (**Figures S1D-L, S2E-H**), broadly confirming the presence of synaptic enhancement in response to temperature increase.

**Figure 1.**
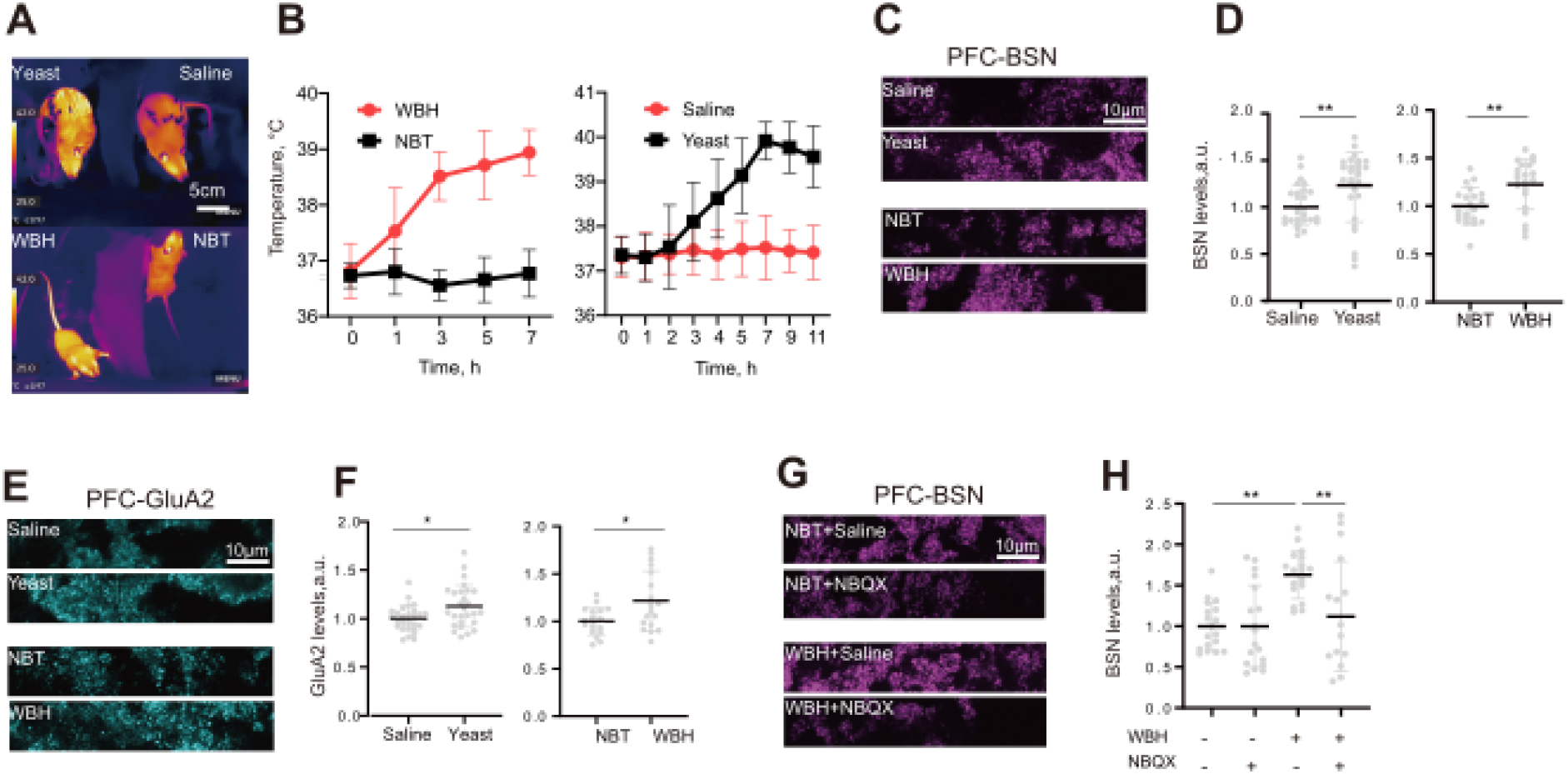
Fever induces synaptic strengthening through AMPARs. (A) Pseudo-colored thermal images showing temperature increase in rats and mouse models. Top, rats, bottom, mice. Left, treated animal, right, control animal. (B) Time course of temperature increase in pyrexia and WBH models and saline controls following protocol induction at t = 0. (C) Images of brain sections from control and treated animals with immunostaining for Bassoon. (D) Quantification of temperature effect on Bassoon levels. **P < 0.01, Student t test; **P < 0.01, Mann-Whitney test. (E) Images of brain sections from control and treated animals with immunostaining for GluA2. (F) Quantification of temperature effect on synaptic GluA2 levels. *P < 0.05, Student t test. (G) Images of brain sections showing the effect of NBQX injection on hyperthermia-induced increase in Bassoon levels. (H) Quantification of the NBQX effect on Bassoon levels. **P < 0.01, Kruskal–Wallis test with Dunn’s multiple comparisons test. N=9/group, pyrexia; 7/group, WBH; 6/group, NBQX+WBH.

Given the key role of glutamatergic neurotransmission in synaptic plasticity, we stained for glutamate receptors, finding an increase in staining for AMPA receptor GluA2 (**Figure 1E-F**), but not NMDA receptor GluN1 (**Figure S2A,B**). Consistent with a role for AMPARs, blockade of AMPARs by a specific antagonist NBQX blocked temperature-induced increase in Bassoon (Figures 1G-H), Homer (Figure S2C,D), Synapsin 1/2 (Figure S2E,F) and Synaptotagmin 1 (Figure S2G,H).. Heat shock proteins Hsp70 and Hsc70 were up-regulated by temperature elevation (**Figure S2I-K**), while intraperitoneal injection of translation inhibitor anisomycin and Hsp70 inhibitor apoptozole both inhibited synaptic enhancement in Homer and Bassoon (**Figure S2L-O**). Thus, AMPAR signaling and heat shock translational response are required for temperature-dependent synaptic enhancement.

In the light of previous evidence linking febrile seizures with long-term alterations in neuronal excitability [1, 2, 7], we next investigated the short- and long-term synaptic effects of juvenile pyrexia. To this end, we induced pyrexia by yeast injection in rats at 4 weeks of age [5], followed by immunohistochemistry at 4 and 10 weeks (**Figures 2Aa, S3A-B**). Pyrexia did not increase Bassoon levels in 4-week animals, indicating the lack of immediate synaptic strengthening in juvenile animals; however, there was significantly more Bassoon puncta in 10-week animals subjected to pyrexia at 4 weeks than in the age-matched controls, indicating long-term synaptic enhancement (**Figure 2B-C**); similar effect was observed for Homer and GluA2 (**Figure S3C-F**), but not for other synaptic markers (**Figure S3G-N**). Levels of Hsp70 and Hsc70 in adult rats were increased at both 4 and 10 weeks, indicating long-term persistence of heat shock response (**Figure S3O-R**).

**Figure 2.**
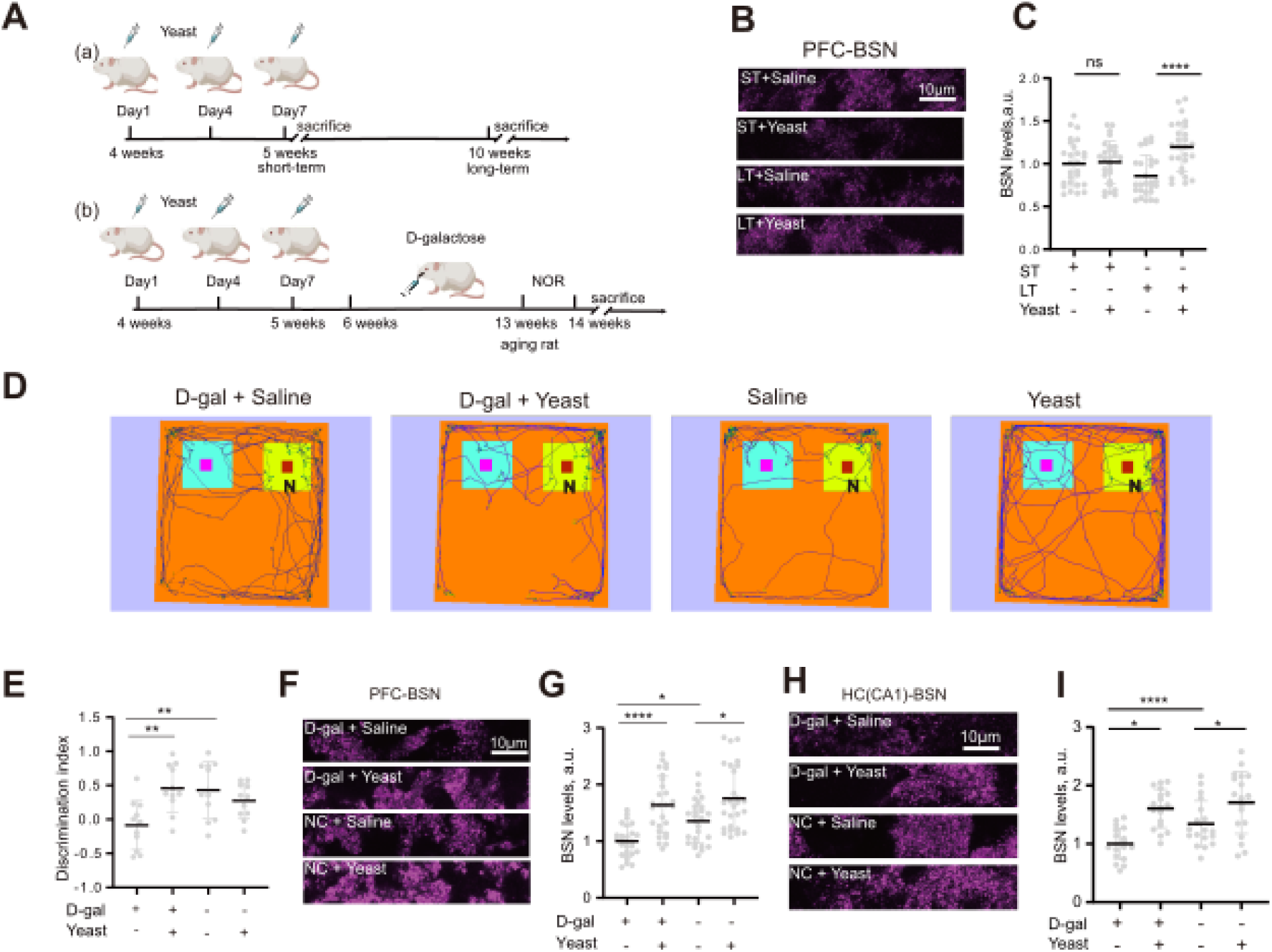
Fever induces long-term synaptic strengthening and protects synapses in a model of aging. (A) Schematics of the experiments for study of the short- and long-term effects(ST and LT) of juvenile pyrexia in rats. (a) - effects of juvenile fever on synapses; (b) - effects of juvenile pyrexia on learning in the model of accelerated aging induced by D-Gal. (B) Images of brain sections showing Bassoon levels immediately after induction of juvenile pyrexia at 4 weeks (short-term, ST) or at 10 weeks following juvenile pyrexia at 4 weeks (long-term, LT). (C) Quantification of Bassoon levels immediately after induction of juvenile pyrexia at 4 weeks (short-term, ST) or at 10 weeks following juvenile pyrexia at 4 weeks (long-term, LT). (D) Representative trajectories during the NOR task. Old object, left (azure); Novel object, right (yellow). (E) Quantification of discrimination indices in control or D-Gal-injected rats with or without prior juvenile febrile experience. **P < 0.01, 1-way ANOVA with multiple comparisons test. (F) Images of brain sections from the prefrontal cortex region (PFC) showing Bassoon levels in control or D-Gal-injected rats with or without prior juvenile febrile experience. (G) Quantification of Bassoon levels from the prefrontal cortex region (PFC) in control or D-Gal-injected rats with or without prior juvenile febrile experience. ****P < 0.0001,*P < 0.05, Kruskal–Wallis test with Dunn’s multiple comparisons test. (H) Images of brain sections from the hippocampal CA1 region (HC-CA1) showing Bassoon levels in control or D-Gal-injected rats with or without prior juvenile febrile experience. (I) Quantification of Bassoon levels from the hippocampal CA1 region (HC-CA1) in control or D-Gal-injected rats with or without prior juvenile febrile experience. ****P < 0.0001,*P < 0.05, 1-way ANOVA with multiple comparisons test. N=9/group, ST and LT pyrexia; 11/group, NOR test; 9/group, PFC staining; 6/group, HC staining.

To test whether juvenile febrile experience can protect against later synaptic dysregulation, we leveraged the D-Gal model of aging (**Figure 2Ab**), which manifests accelerated synaptic loss and memory decline [8]. We then used the novel object recognition test (NOR) to assess the effects of D-Gal and juvenile pyrexia experience on spatial learning. At 13 weeks, total observation time and discrimination index were significantly lower in D-Gal-treated rats than in controls (**Figures 2D-E, S4A**). This cognitive defect was mirrored by a decrease in Bassoon and Homer intensity as measured by immunostaining in the PFC and the hippocampal CA1 region at 14 weeks, indicative of synaptic loss accompanying learning deficits (**Figures 2F-I, S4D-G**). Conversely, animals with a prior experience of pyrexia at 4 weeks of age showed no D-Gal-induced decline in NOR and synaptic staining, suggesting that juvenile fever had a protective effect on synapses and preserved spatial learning ability (**Figures 2D-I, S4A, S4C, S4D-G**). There were no significant differences in total travelled distances between experimental groups (**Figure S4B**), indicating that overall mobility was not affected by treatments.

Our study demonstrates that moderate fever is sufficient to trigger long-term enhancement of PFC synapses *in vivo*, safeguarding cognition in a model of aging. AMPAR- and translation-dependent synaptic enhancement is consistent with homeostatic scaling [9] which may in turn be linked to systemic activity patterns *e.g.* hyperthermia-induced spreading depressions [10]. While mechanistic understanding of fever-induced synaptic remodelling requires an in-depth study, possible mechanisms underlying this phenomenon may include hyperthermia-induced BDNF release [11, 12] and direct enhancement of AMPAR activity via glutamate gating [13]. At this preliminary stage, our study focused on males to avoid the confounding cause-effect relationship between temperature and the estrous cycle [14, 15] and employed an accelerated aging model. Further exploration of the putative therapeutic potential of fever for synaptic protection will necessitate systemic investigation of pyrexia-associated synaptic remodelling in the context of natural aging in both sexes, with a particular focus on age-related synaptopathies, as well as detailed safety profiling in the clinic [16].

## Materials & Methods

### Reagents, drugs, fluorescent markers

Institute for Cancer Research (ICR) mice and Sprague-Dawley rats were from Jinan Pengyue Laboratory Animal Co. Apoptozole and NBQX were from AbMole. Anisomycin and D-galactose were from Macklin. Yeast was from Angle Yeast. Antibodies against gephyrin, vGlut1, synapsin 1/2, GABRA1, Cav2.1, synaptotagmin 1 and Homer were from Synaptic Systems. BSN was from Abcam. Antibody against GluA2 was from Merck Millipore. Antibody against GluN1 was from Alomone Labs. Antibodies against Hsp70 and Hsc70 were from Boster Bio. Rabbit IgG conjugated to AlexaFluor488 was from Bioss. Mouse IgG conjugated to AlexaFluor594 was from SolarBio.

### Animals

Male, 8-10 week-old, 28-32 g of weight, specific pathogen free (SPF), Institute of Cancer Research (ICR) mice were used in the whole-body hyperthermia model. Male, 8-10 week-old, 230-250 g of weight, SPF Sprague Dawley (SD) rats were used in the yeast-induced pyrexia model. Male, 3-4 week-old, 60-80 g of weight, SPF SD rats were used in the juvenile pyrexia model. All animal use and experimental protocols were approved and carried out in compliance with the IACUC guidelines and the Animal Care and Ethics Committee of Qingdao University School of Medicine (approval number QDU-HEC-2024458).

All the animals were housed in the animal facility of Qingdao University, under conditions adhering to the GB 14925-2010 standard of Laboratory Animal Environment and Facilities. Specifically, the temperature was maintained between 18-26°C, with a day-night temperature difference not exceeding 4°C. The relative humidity was kept at 40-70% to prevent proliferation of pathogens. Ventilation system provided 10-20 air exchanges per hour, with ammonia concentrations not exceeding 14 mg/m^3^. The lighting cycle was set at 12 hours of light followed by 12 hours of darkness, with a lighting intensity of 150-300 lx. Noise levels were controlled to maintain the animals’ normal physiological rhythms and reduce stress responses. Animal rooms were maintained at the SPF-grade cleanliness standard.

### Whole body hyperthermia model

Animals were randomly allocated into WBH and normal body temperature (NBT). The WBH group was put into a heating box, and the temperature was increased by 0.5°C every 15 min until it reached 39°C. Temperature was maintained at 39°C for 6 hours, with a rest every 2 hours in which all mice were placed at room temperature for 15 minutes while rectal temperature was measured and 0.1 mL saline was injected intraperitoneally, before returning to the 39°C environment. Rectal temperature was measured at a final rest immediately before thermal imaging. NBT was maintained at room temperature (18-22°C), with the same procedures for injection and temperature measurement as described above. All animals were sacrificed one hour later.

### Yeast-induced fever model

Animals were divided into two groups: pyrexia (Yeast) and control (Saline only). Yeast was dissolved in saline at 37°C and adjusted to a 20% yeast solution. The rats in the Yeast group received subcutaneous injection of 20% yeast solution to a final concentration of 10 mL/kg body weight in the posterior neck, and the rats in the Saline group received subcutaneous injection of the same volume of normal saline. Rectal temperature was measured at the beginning of the injection and at one- or two-hour intervals thereafter, thermal imaging was performed at the seventh hour after injection, and animals were sacrificed at the twelfth hour after injection.

### Juvenile fever model

Animals were divided into two groups as above. For juvenile pyrexia induction, animals were subcutaneously injected with 20% yeast 10 ml/kg at the posterior neck. Injections were given every 2 days for a total of 3 injections in a week (on the first, fourth, and seventh days of the week). Rectal temperature measurement was also performed at a fixed time during the week of injection to monitor body temperature changes. The control group was injected with the same volume of saline at the same time and rectal temperature was measured. Thermal imaging was performed 7 h after the third injection. The animals were sacrificed at 12 h after the last injection of the current week or after 6 weeks of rearing (approximately 10 weeks).

### Injection of inhibitors

NBQX, anisomycin and apoptozole were injected intraperitoneally one hour before the whole body hyperthermia model. The control group was given the same amount of normal saline at the concentrations of 10 mg/kg, 150 mg/kg and 4 mg/kg respectively. The intraperitoneal injection volume was 100 μl per animal.

### Accelerated aging model

At 6 weeks, animals previously subjected to juvenile fever were given D-galactose 300 mg/ml/kg/d by gavage in water (09:00-10:00 am) for 7 weeks (about 13 weeks of age). The control group was given the same amount of water by gavage. At 14 weeks of age, animals were subjected to behavioral experiments for novel object recognition experiments, and animals were sacrificed after 14 weeks.

### Novel object recognition testing

At 13 weeks of age, environmental exposure was carried out at the same time of a day over a 7-day period. Animals were randomly assigned and blinded to the experimenter to avoid bias. Three days before the experiment, animals were placed in the open field box to explore freely for 10 min to adapt to the environment. Placing two identical objects, A and B, symmetrically in the test box on the day of the experiment ensured that the objects were not pushed. The distance between two identical objects was 60 cm and 20 cm, respectively. Animals were placed in the experimental box with their back to the subject object and equidistant from the subject object. Familiarization period with the objects lasted for 5 minutes, during which time animals were allowed to freely explore the objects within the testing arena. For 40 minutes after the familiarization phase, animals were returned to their home cages to facilitate memory consolidation. This was followed by the 10-minute long test session, during which one of the familiar objects was replaced with a novel one, and the animals’ exploratory behavior was recorded to evaluate their recognition memory. The number, time and distance of exploration within 10 min were recorded. The video data was used to calculate the proportion of time the rat spent exploring the old objects (F) and novel objects (N).

Recognition Index (RI) was calculated as

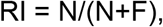

Discrimination Index was calculated as

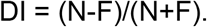

Difference in recognition bouts was calculated as the number of exploration bouts for novel objects divided by the sum of exploration bouts for novel and old objects.

### Immunohistochemistry staining

After treatment, animals were anesthetized with isoflurane and then perfused transcardially with normal saline. The brains were dissected out and kept at -80°C. Coronal sections (25 um) were cut at -20°C with a cryostat (Leica CM1950) Sections were made at approximately +1.5 to +3, +1.5 to +3 and +2 to +4 mm from bregma in mice, juvenile rats and adult rats respectively.

then fixed with 4% PFA for 20 mins at RT and washed three times for 10 mins each in PBS. Sections were incubated in blocking buffer (10% goat serum, 0.3% Triton-X 100 in PBS) for 1 hour at RT, and incubated with primary antibodies for 36-48h at 4°C, followed by 1 hour at RT. Sections were washed six times for 5 mins each in PBS, followed by incubating with Alexa Fluor 488- and Alexa Fluor 594-conjugated secondary antibodies for 75 mins at room temperature, and washed three times for 10 mins each in PBS. Then all sections were mounted onto positively-charged microscope slides and covered in DAPI-containing mount media (Solarbio). To control for specificity of staining, negative controls were conducted with only secondary antibodies, showing that the signal intensity due to non-specific binding was <5%.

### Confocal microscopy

Samples were imaged on Nikon Eclipse Ti2 laser confocal microscope. The imaging systems were controlled by NIS Elements 2.0 software. The imaging parameters for Nikon Eclipse Ti2 were as follows. The following image acquisition settings were used for serial confocal z-stack images: tissue sections of synaptic proteins - 0.5 μm step, range: 2 μm, 512 × 512 pixels, 2x zoom, 100x magnification; tissue sections of HSP -512 × 512 pixels, 1x zoom, 20x magnification. Pinhole size was kept at a suitable (1-2AU) Airy unit. Excitation laser wavelengths were 488 nm and 561 nm. Bandpass filters were set at 500−550 nm (FITC, Alexa Fluor 488) and 570−620 nm (TRITC, AlexaFluor 594). Gain, exposure and offset settings were optimized within each experiment to ensure appropriate dynamic range, low background and optimal signal/noise ratio.

### Image analysis

To identify individual synapses, images were binarized in ImageJ using the”Moments” setting, and particles were counted automatically using the “Analyze Particles”command across the whole image. Signal intensities were quantified for each synaptic punctum using the Region of Interest (ROI) Manager function of ImageJ. To avoid rare overlap of multiple synapses, only ROIs with areas ranging from 0.1 to 2 μm^2^ were included in further analysis. All values of circularity were included in analysis. Individual ROIs were then combined into one compound ROI using the “Combine” and “Add” functions of the ROI Manager interface, whereupon quantification of mean signal intensity in each channel was performed using the “Measure” function. Since background fluorescence intensity was typically less than 1% of the median fluorescence in each channel within a ROI, background subtraction did not significantly affect the measurements and was not performed.

## Statistical analysis

All the experiments were performed in at least 6 independent biological replicates, with three fields of view from one brain section per animal used for analysis. Statistical analysis was carried out using the GraphPad Prism v5 (https://www.graphpad.com/). For each experiment, data was normalised to the average value in the control sample. Data distributions were assessed for normality using d’Agostino and Pearson omnibus normality tests. For normally distributed datasets, two sample t test and 1-way ANOVA, with Dunn’s, Bonferroni’s post tests were used to assess statistical significance as appropriate; for non-normally distributed datasets, Mann-Whitney rank test, Kruskal-Wallis test and Dunn’s post test were used. The data sets are presented in the form of scatter plots and line plots, with plot averages as lines.

## Sex as a biological variable

Male animals were used throughout the study.

## Study approval

All animal use and experimental protocols were approved and carried out in compliance with the IACUC guidelines and the Animal Care and Ethics Committee of Qingdao University School of Medicine.

## Data availability

Raw experimental data is available from the Authors upon written request.

## Acknowledgements

This work was funded by the Lewy Body Society (OOG2019/2020) and the National Natural Science Foundation of China (32070772).

## Supplementary Figures

**Figure S1.**
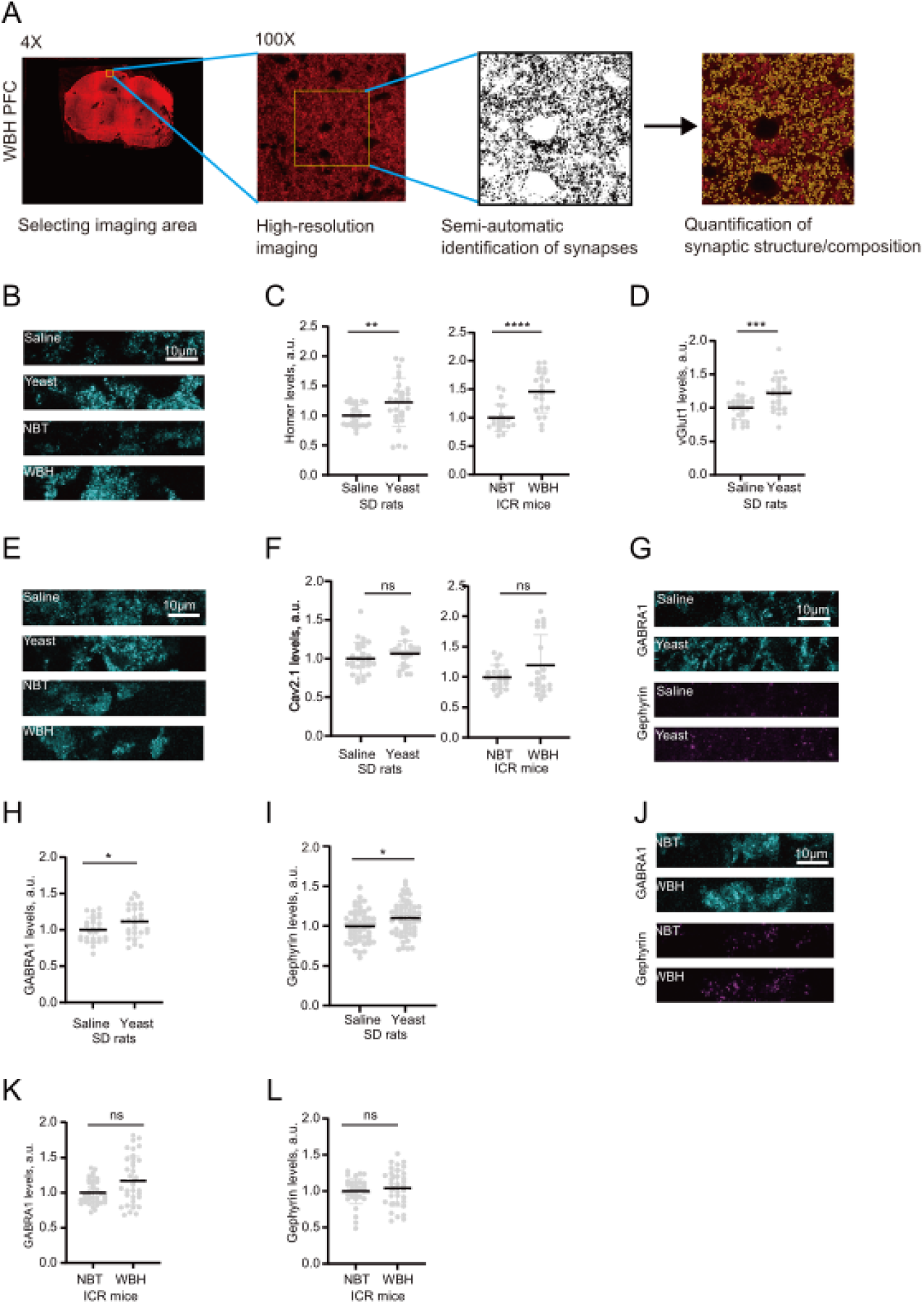
Temperature enhances various synaptic markers. (A) Schematic of the experimental approach to quantification of synaptic structure and composition. For details, see Materials and Methods, sections “Confocal Microscopy” and “Image Analysis”. (B) Images of brain sections from control and treated animals with immunostaining for Homer. (C) Quantification of temperature effect on Homer levels in control and treated animals. (D) Quantification of temperature effect on vGlut1 levels in control and treated animals. (E) Images of brain sections from control and treated animals with immunostaining for CaV2.1. (F) Quantification of temperature effect on CaV2.1 levels in control and treated animals. (G) Images of brain sections from control and yeast-injected animals with immunostaining for GABRA1& gephyrin. (H) Quantification of temperature effect on GABRA1 levels in control and yeast-injected animals. (I) Quantification of temperature effect on gephyrin levels control and yeast-injected animals. (J) Images of brain sections from control and WBH-treated animals with immunostaining for GABRA1& gephyrin. (K) Quantification of temperature effect on GABRA1 levels in control and WBH-treated animals. (L) Quantification of temperature effect on gephyrin levels control and WBH-treated animals. ****P<0.0001, ***P<0.001, **P<0.01, *P<0.05, ns - not significant. N=9/group, pyrexia; 7/group, WBH.

**Figure S2.**
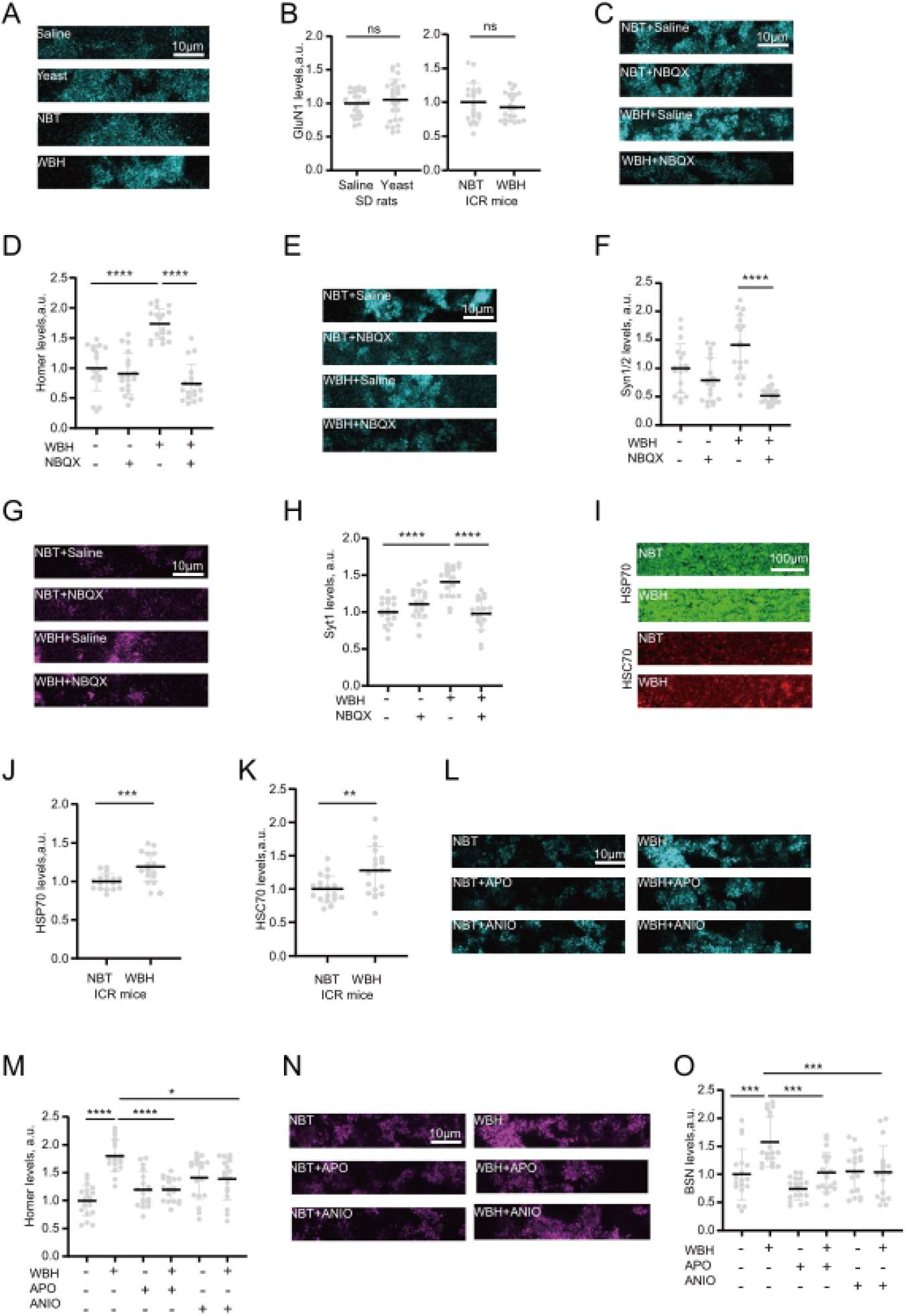
Mechanisms of synaptic potentiation induced by fever. (A) Images of brain sections from control and treated animals with immunostaining for GluN1. (B) Quantification of temperature effect on GluN1 levels in control and treated animals. (C) Images of brain sections from control and WBH-treated animals showing NBQX effect on WBH-induced increase in Homer. (D) Quantification of NBQX effect on WBH-induced increase in Homer. (E) Images of brain sections from control and WBH-treated animals showing WBH-induced increase in synapsin 1/2 and NBQX effect thereon. (F) Quantification of WBH-induced increase in synapsin 1/2 and NBQX effect thereon. (G) Images of brain sections from control and WBH-treated animals showing WBH--induced increase in synaptotagmin 1 and NBQX effect thereon. (H) Quantification of WBH-induced increase in synaptotagmin 1 and NBQX effect thereon. (I) Images of brain sections from control and WBH-treated animals showing WBH-induced increase in Hsp70 and Hsc70. (I) Quantification of WBH-induced increase in Hsp70. (K) Quantification of WBH-induced increase in Hsc70. (L) Images of brain section from control and treated animals showing effects of apoptozole and anisomycin application on temperature-induced Homer levels. (M) Quantification of the effects of apoptozole and anisomycin application on temperature-induced Homer levels. (N) Images of brain section from control and treated animals showing effects of apoptozole and anisomycin application on temperature-induced Bassoon levels. (O) Quantification of the effects of apoptozole and anisomycin application on temperature-induced Bassoon levels. ****P<0.0001, ***P<0.001, **P<0.01, *P<0.05, ns - not significant. N≥6 independent biological replicates. N=9/group, pyrexia; 7/group, WBH; 6/group, NBQX+WBH; 6/group, Anisomycin+apoptozole.

**Figure S3.**
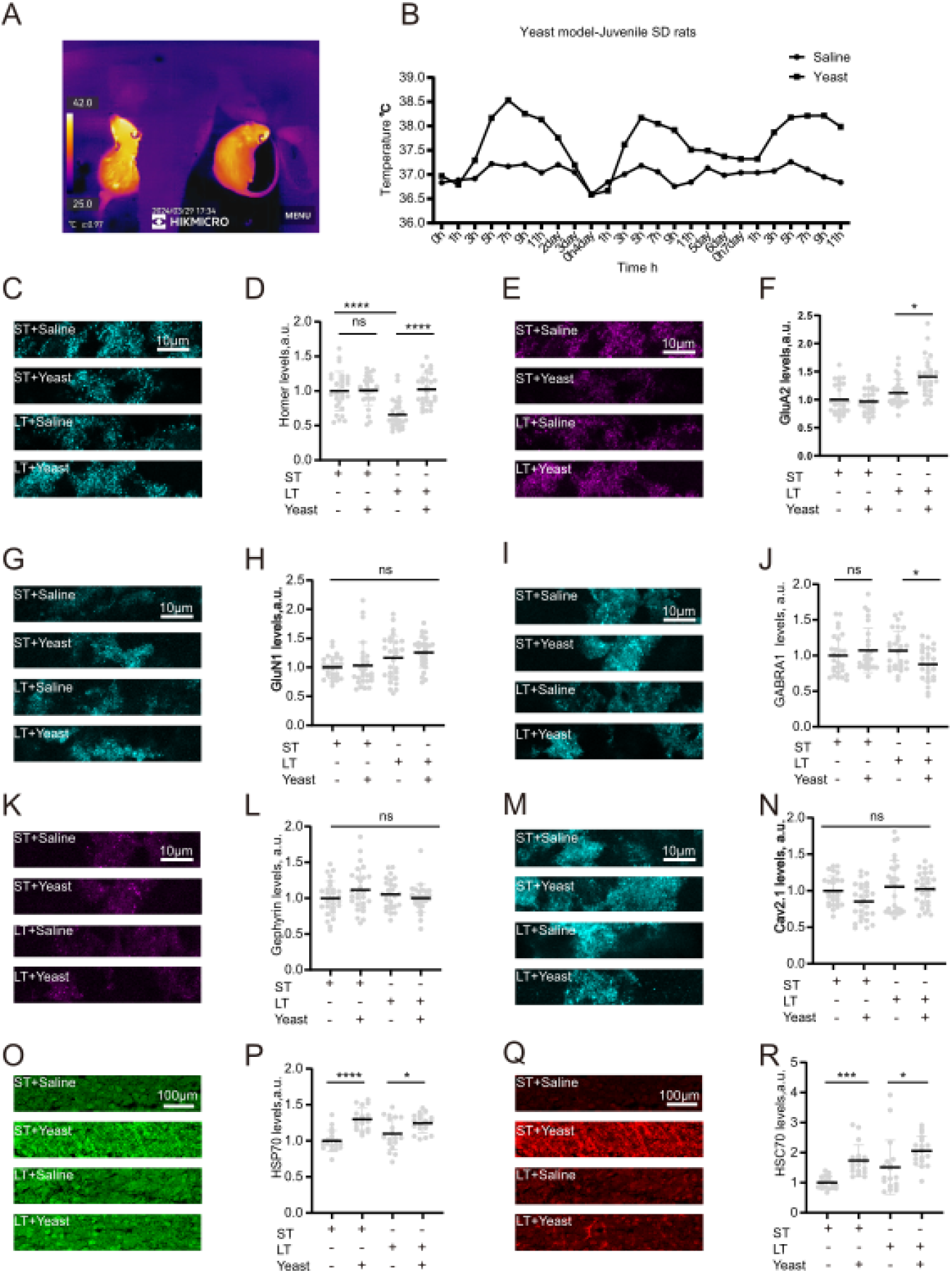
Short- and long-term synaptic changes following juvenile pyrexia. (A) Images of a 4-week old rat subjected to pyrexia induction by yeast injection (left) and a control rat with saline injection (right). (B) Example temperature traces showing repeated bouts of temperature elevation in the juvenile fever model. (C) Images of brain sections showing the effect of juvenile pyrexia on Homer levels at 4 (short-term, ST) and 10 (long-term, LT) weeks. (D) Quantification of the juvenile pyrexia effect on Homer levels at 4 (short-term, ST) and 10 (LT) weeks. N=9. (E) Images of brain sections showing the effect of juvenile pyrexia on GluA2 levels at 4 (short-term, ST) and 10 (long-term, LT) weeks. (F) Quantification of the juvenile pyrexia effect on GluA2 levels at 4 (short-term, ST) and 10 (long-term, LT) weeks. (G) Images of brain sections showing the effect of juvenile pyrexia on GluN1 levels at 4 (short-term, ST) and 10 (long-term, LT) weeks. (H) Quantification of the juvenile fever effect on GluN1 levels at 4 (short-term, ST) and 10 (LT) weeks. (I) Images of brain sections showing the effect of juvenile pyrexia on GABRA1 levels at 4 (short-term, ST) and 10 (long-term, LT) weeks. (J) Quantification of the juvenile pyrexia effect on GABRA1 levels at 4 (short-term, ST) and 10 (long-term, LT) weeks. (K) Images of brain sections showing the effect of juvenile pyrexia on gephyrin levels at 4 (short-term, ST) and 10 (long-term, LT) weeks. (L) Quantification of the juvenile pyrexia effect on gephyrin levels at 4 (short-term, ST) and 10 (long-term, LT) weeks. (M) Images of brain sections showing the effect of juvenile pyrexia on CaV2.1 levels at 4 (short-term, ST) and 10 (long-term, LT) weeks. (N) Quantification of the juvenile pyrexia effect on CaV2.1 levels at 4 (short-term, ST) and 10 (long-term, LT) weeks. (O) Images of brain sections showing the effect of juvenile pyrexia on Hsp70 levels at 4 (short-term, ST) and 10 (long-term, LT) weeks. (P) Quantification of the juvenile pyrexia effect on Hsp70 levels at 4 (short-term, ST) and 10 (long-term, LT) weeks. (Q) Images of brain sections showing the effect of juvenile pyrexia on Hsc70 levels at 4 (short-term, ST) and 10 (long-term, LT) weeks. (R) Quantification of the juvenile pyrexia effect on Hsc70 levels at 4 (short-term, ST) and 10 (long-term, LT) weeks. ns-not significant, *P < 0.05, ***P < 0.001,****P < 0.0001, Kruskal-Wallis test with Dunn’s multiple comparisons test. N=9/group, ST and LT pyrexia.

**Fig. S4.**
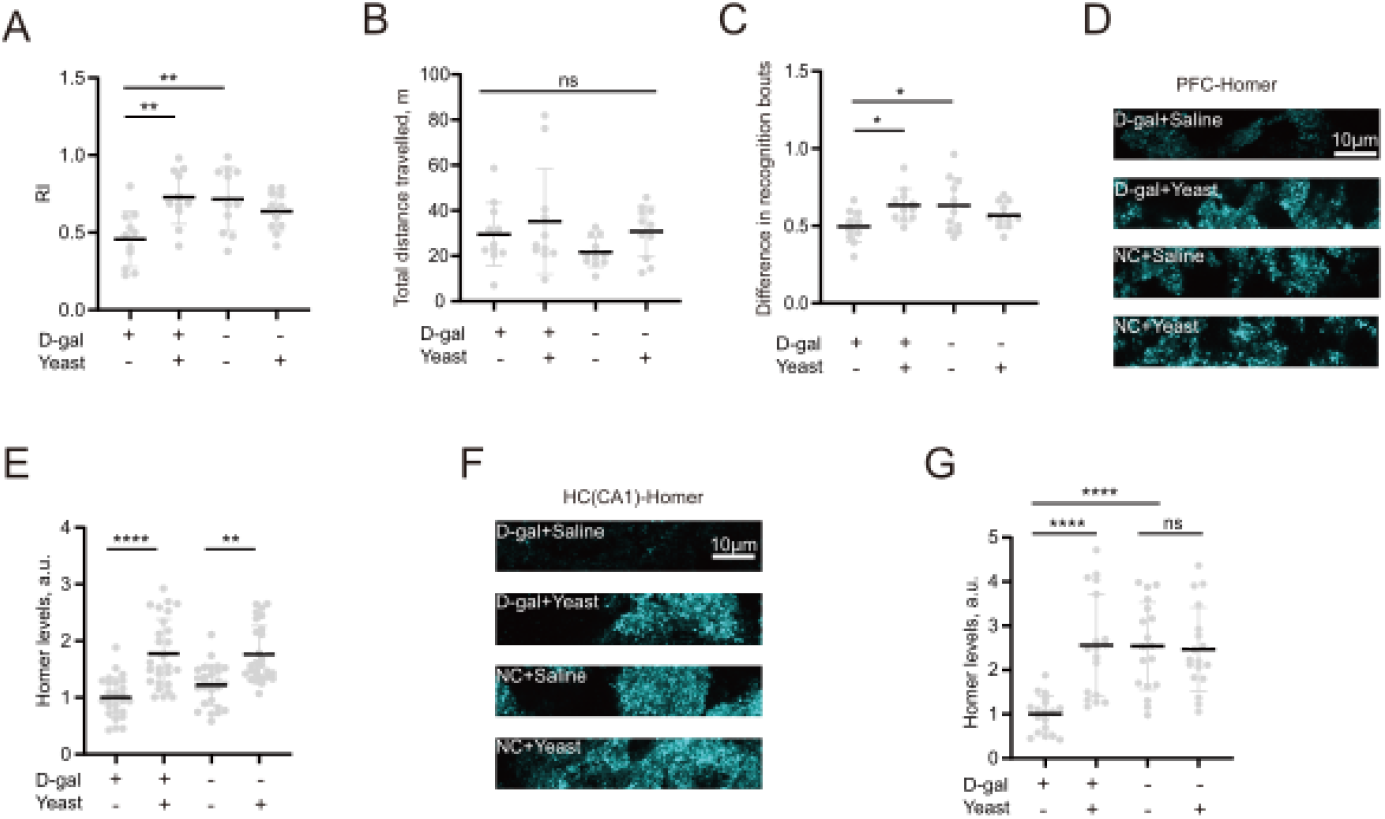
Long-term effects of juvenile pyrexia in the D-Gal injected rats. (A) Effects of juvenile pyrexia on spatial preferences in control and D-Gal-injected rats with or without prior juvenile febrile experience. (B) Effects of juvenile pyrexia on total distance in control and D-Gal-injected rats with or without prior juvenile febrile experience. (C) Effects of juvenile pyrexia on Recognition Bouts in control and D-Gal-injected rats with or without prior juvenile febrile experience. (D) Images of brain sections from the prefrontal cortex region (PFC) showing Homer levels in control or D-Gal-injected rats with or without prior juvenile pyrexia. (E) Quantification of Homer levels from the prefrontal cortex region (PFC) in control or D-Gal-injected rats with or without prior juvenile pyrexia. ns-not significant, *P < 0.05, **P < 0.01,****P < 0.0001, Kruskal-Wallis test with Dunn’s multiple comparisons test. (F) Images of brain sections from the hippocampal CA1 region (HC-CA1) showing Homer levels in control or D-Gal-injected rats with or without prior juvenile pyrexia. (G) Quantification of Homer levels from the hippocampal CA1 region (HC-CA1) in control or D-Gal-injected rats with or without prior juvenile pyrexia. ****P < 0.0001, ns-not significant,, 1-way ANOVA with multiple comparisons test. N=11/group, NOR test; 9/group, PFC staining; 6/group, HC staining.

